# Population genomic analysis of the speckled dace species complex (*Rhinichthys osculus*) identifies three species-level lineages in California

**DOI:** 10.1101/2021.12.14.472667

**Authors:** Yingxin Su, Peter B. Moyle, Matthew A. Campbell, Amanda J. Finger, Sean M. O’Rourke, Jason Baumsteiger, Michael R. Miller

**Affiliations:** Department of Animal Science, University of California, Davis, California, United States of America; Department of Wildlife, Fish, and Conservation Biology, University of California, Davis, California, United States of America; Center for Watershed Sciences, University of California, Davis, California, United States of America

## Abstract

The speckled dace (*Rhinichthys osculus*) is small cyprinid fish that is widespread in the Western USA. Currently treated as a single species, speckled dace consists of multiple evolutionary lineages that can be recognized as species and subspecies throughout its range. Recognition of taxonomic distinctiveness of speckled dace populations is important for developing conservation strategies. In this study, we collected samples of speckled dace from 38 locations in the American West, with a focus on California. We used RAD sequencing to extract thousands of SNPs across the genome from samples to identify genetic differences among seven California populations informally recognized as speckled dace subspecies: Amargosa, Owens, Long Valley, Lahontan, Klamath, Sacramento, and Santa Ana speckled dace. We performed principal component analysis, admixture analysis, estimated pairwise Fst, and constructed a phylogeny to explore taxonomic relationships among these groups and test if these subspecies warrant formal recognition. Our analyses show that the seven subspecies fit into three major lineages equivalent to species: western (Sacramento-Klamath), Santa Ana, and Lahontan speckled dace. Death Valley speckled dace were determined to be two lineages (Amargosa and Long Valley) within Lahontan speckled dace. Western and Lahontan speckled dace lineages had branches that can be designated as subspecies. These designations fit well with the geologic history of the region which has promoted long isolation of populations. This study highlights the importance of genetic analysis for conservation and management of freshwater fishes.

## Introduction

The speckled dace (*Rhinichthys osculus*) is a small (usually <10 cm total length) cyprinid fish that is widely distributed across western North America. It is found from northern Mexico and southern California through central and northern California, the Great Basin, the Pacific Northwest, to southwestern Canada [1-3]. Despite its wide distribution, the speckled dace is considered to be one highly variable species, albeit with numerous subspecies, many of which are undescribed [3]. Here we refer to the species as the speckled dace complex (SDC). The SDC diverged from the longnose dace *R. cataractae* species complex of eastern North America over 6 million years ago [4]. The common ancestor of the SDC was presumably initially isolated in the ancestral waterway of the Columbia River and then spread throughout the western USA and British Columbia as the result of geologic events that connected and disconnected watersheds [3]. Populations are found in a wide array of habitats, from desert springs to large rivers and lakes, but most typically to small to medium-sized streams. Their morphology is highly variable but generally reflects the habitat in which a particular population lives. For example, narrow caudal peduncles and large pectoral fins characterize swift-water populations and more robust bodies, thicker caudal peduncles and smaller pectoral fins characterize quiet-water populations [3,5,6].

Historically, many populations of the SDC were described as separate species. Jordan and Evermann (1896) list 10 species, which had mostly been described based partially on their isolation from other populations and partially on morphological and meristic characteristics even though these characters overlapped among populations [7]. The species were all placed in the genus *Apocope* (now a subgenus of *Rhinichthys*). Subsequently, most of these forms were united under *R. osculus* and considered at best to be subspecies [8]. However, the presence of many isolated populations of speckled dace with similar adaptations to local environments and hence convergent morphologies suggests that cryptic species [9] exist within the SDC and that some of the recognized subspecies (listed in [3]) could be considered as species.

In California, based on the early taxonomic literature descriptions of life history traits, and co-occurrence in isolated basins with other endemic fishes, Moyle (2002) recognized speckled dace as one species with seven subspecies: Lahontan speckled dace (*R. o. robustus*), Klamath speckled dace (*R. o. klamathensis)*, Sacramento speckled dace (*R. o. subsp*.), Owens speckled dace (*R. o. subsp*.), Long Valley speckled dace (*R. o. subsp*.), Amargosa speckled dace (*R. o. nevadensis*), and Santa Ana speckled dace (*R. o. subsp*.) [1]. Differences in morphology and meristics among these subspecies are small and may reflect local adaptations rather than fixed characteristics (Smith et al. 2017).

The advent of molecular genetic techniques has resulted in renewed efforts to examine diversity within the SDC. Genetic information is used to develop hypotheses of evolutionary relationships among populations and to generate biogeographic scenarios relating speckled dace to the history of the western aquatic landscape [3]. To date, the primary genetic approach used to investigate the systematics of speckled dace was analyses of mitochondrial DNA [3,10]. Oakey et al. (2004) used 112 restriction sites found in the mitochondrial genome of dace distributed across the western USA to construct a molecular phylogeny of the speckled dace species complex [10] and found a close match between MtDNA patterns and the geologic history and isolation of drainage basins. They concluded that the SDC consisted of three main evolutionary lineages [9]: (1) Colorado River Basin and Los Angeles Basin, (2) Great Basin (Snake River, Bonneville, Death Valley and Lahontan) and (3) Columbia and Klamath-Pit Rivers. Pfrender et al (2004) showed that MtDNA patterns reflected long isolation of populations in five river basins in Oregon and suggested that some of the lineages were distinct enough to be considered species [11]. In contrast, Billman et al (2010) did not find species-level differences in MtDNA among SDC from Great Basin waterways (Snake, Bonneville, Lahontan) [12].

More narrowly, Ardren et al. (2010) applied MtDNA analysis to the Foskett Spring speckled dace population in the Warner Basin, Oregon [13]. They concluded that this dace was not sufficiently different from other dace to be considered even a subspecies, although members of the SDC from throughout the Warner Basin together were distinct at the species level. Hoekzema and Sidlauskas (2014) also examined SDC fish from the Warner Basin, along with dace from five other isolated populations in the Great Basin in Oregon [14]. They used MtDNA as well as nuclear DNA (nuclear s7 intron) and found that dace in the Warner Basin were different, potentially at the species level, from dace in the other four basins.

The most comprehensive study of the SDC was that of Smith et al. (2017) who compared dace populations from throughout western North America, using MtDNA, morphology, fossils, and the geologic record of the entire region. While their analyses indicated multiple lineages, they concluded that there was considerable, if sporadic, gene flow among populations, reflecting complex geologic events that promoted both connectivity and isolation. According to their analysis, gene flow prevented the formation of morphologically distinct populations that might be defined as species, through the process of reticulate evolution.

Recognizing the limitations of MtDNA for determining evolutionary lineages, Mussman et al. (2020) compared populations of speckled dace from throughout the Death Valley region, in the Owens and Amargosa river basins, using double-digest RAD [15]. They found that Death Valley has four distinct evolutionary lineages that they placed in one subspecies of *R. osculus*: the Amargosa speckled dace (*R*.*o. nevadensis*) with each of the lineages being treated as a Distinct Population Segment (DPS) for management purposes [15].

Overall, genetic studies have produced mixed results as to whether or not any evolutionary lineages in the SDC are distinct enough to be designated as species or subspecies. The default position is to follow Smith and Dowling (2008) and Smith et al. (2017) that the SDC is a single species throughout its range because the various populations lack unique morphometric characteristics that would allow them to be described as species [3-4]. However, this default position is particularly problematic for California, a region rich in endemic fish species, many of which are threatened with extinction [1,16]. California SDC populations are also among those most distant from the hypothesized region of origin in the Columbia River and are among the most southern of the taxon. These SDC populations thus reflect their remarkable record of colonizing new regions during the wetter periods of the Pleistocene and then adapting to new conditions as areas became drier [3] and more isolated.

In this paper, we analyze speckled dace relationships using genomics, more specifically, restriction-site associated DNA sequencing (RAD-seq). This approach is well suited for analyzing the SDC because it uses thousands of loci distributed across the genome from each individual rather than only a single locus or handful of loci as was possible with previous methods. For further discussion of this approach to resolving issues with identifying cryptic fish species, see Baumsteiger et al. (2017) [17].

We investigated the following questions using a genome-wide data set: (1) Is the SDC just one species or multiple species throughout its native range but especially in California? (2) Are the subspecies of speckled dace found in California, as listed in Moyle (2002), supported by genomic analysis? (3) Do analyses of the SDC using genomic techniques confirm evolutionary relationships inferred from other methods of analysis, especially use of MtDNA? (4) Can we designate species and subspecies of speckled dace base on genetic distinctiveness, monophyly, and geographical isolation?

## Methods

### 1.1 Sampling and DNA sequencing

To delineate the lineages of SDC, we obtained samples from 38 locations across the major zoogeographic regions (Fig 1 & S1 Appendix). Fin clips were taken from live adults or from whole fish stored in ethanol and dried on Whatman qualitative filter paper and stored at room temperature. DNA was extracted from fin clips with a magnetic bead–based protocol [18] and quantified using Quant-iT PicoGreen dsDNA Reagent (Thermo Fisher Scientific) with an FLx800 Fluorescence Reader (BioTek Instruments). Genomic DNA was used to generate *SbfI* RAD libraries [18] and sequenced with paired-end 100-bp reads on an Illumina HiSeq 2500. Demultiplexing was performed requiring an exact match with well and plate barcodes [18]. Sequencing coverage was assessed at the 50 bp position of each *de novo* RAD contig (see below) across all individuals using the depth function in SAMtools [19].

**Fig 1.**
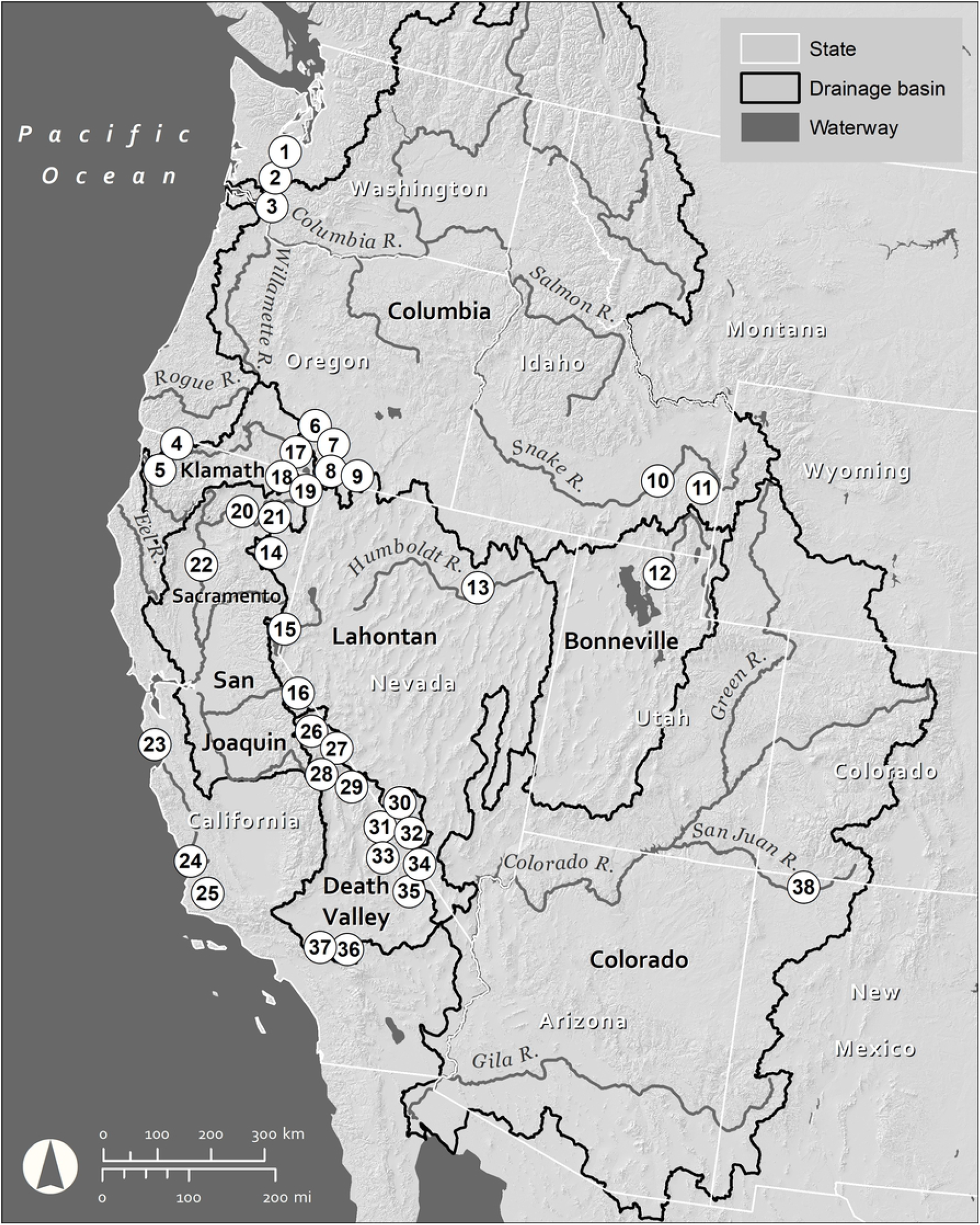
Sampling map. Map of sampling sites in which speckled dace were collected in this study. The location represented by each number can be found in the supplementary material, Appendix 1.

### 1.2 RAD De Novo Assembly and Alignments

To generate a reference sequence for speckled dace, we performed RAD *de novo* assembly on 8 individuals from the Walker River (S1 material). Specific details of the *de novo* assembly methods may be found in Baumsteiger et al. (2017), but, briefly, a bioinformatic pipeline including a genome assembler was used to construct a partial reference for speckled dace [17]. After *de novo* assembly, the mem in the Burrows–Wheeler aligner (BWA) was used to align each sample to the reference under the default parameters. SAMtools was used to convert SAM files to BAM files, calculate the percentage of aligned reads, remove PCR duplicates, filter for the proper pairs, and merge the alignments if needed [19]. After the removal of PCR duplicates, we removed low-coverage individuals with less than 70,000 mapped reads.

### 1.3 Genetic population structure with PCA

To begin investigating population structure, we used Analysis of Next Generation Sequencing Data (ANGSD) to call SNPs (-SNP_eval1e-12), infer major and minor alleles (-doMajorMinor 1), and estimate allele frequencies (-doMaf 2). Only reads with a mapping quality score above 20 (-minMapQ 20) and only bases with a quality score above 20 (-minQ 20) were used in this process [20]. Furthermore, only SNPs with a minor allele frequency of at least 0.01 (-minMaf) that were represented in at least 50% of the included samples (-minInd 88). These SNPs were then used to calculate a covariance matrix (-doCov 1), which was used to generate eigenvalues and eigenvectors for Principal Component Analysis (PCA). The percentage of total genetic variation explained by each PC was calculated, and PCs explaining a relatively large proportion of genetic variation were plotted with ggplot2. To view the substructures within groups from the initial PCA, subsequent PCAs were performed on samples from each group using the same methods described above. The number of SNPs used in each substructure analysis is listed in S1 Table.

### 1.4 Genetic population structure with Admixture analysis

To further assess population structure in speckled dace, we generated genotype likelihoods with ANGSD using the same parameters as above. The beagle output file was then used as the input file for NGSadmix [21]. The parameter K, which means the number of clusters that samples are partitioned assumed in each analysis, were run from 2 to 9, and each has a minor allele frequency filter of 0.01. After population structure was initially characterized, we repeated the procedure as described above on subsets of samples to investigate substructure within each group.

### 1.5 Quantifying pairwise divergence between genetic lineages

To quantify the genetic divergence among populations, we calculated genome-wide F_st_ for population units identified by the analysis above. The individual BAM files were grouped by seven subspecies designated in Moyle (2002) and undesignated speckled dace were grouped by the geographical range. The folded site allele frequencies (SAF) were estimated for each group. The SAF file for the pairwise locations were the input to estimate two-dimensional site frequency spectrum (SFS). SAF for each location and two-dimensional SFS were used to estimate weighted genome-wide F_st_. All the steps are processed in RealSFS set in ANGSD.

### 1.6 Molecular Phylogeny

To further investigate the relationships among different genetic lineages, a range-wide phylogenetic tree was generated using SVDQuartets [22-23]. Relict dace (*Relictus solitarius*) and tui chub (*Siphatales bicolor*) were used as the outgroup to root the molecular phylogeny because both species are western cyprinids (material S2). The tips were assigned to the subspecies described in Moyle (2002). Undesignated speckled dace were represented by the location where they were collected. If a significant genetic difference was shown between the locations in one region or between subspecies in PCA or admixture analysis, the group was separated into two tips based on genetic differences shown in the other analysis.

We used ANGSD to perform genotype calling, and we used the same parameters as mentioned above, except generating a VCF file (-dovcf 1). BCFTOOLS were used to prune the SNPs with r^2^ greater than 0.9 within each RAD contig [24]. The pruned VCF file was transformed into NEXUS format by vcf2phylip [25]. The pruned NEXUS file was analyzed by SVDQuartets loaded within PAUP° 4.0 [26]. We selected multispecies coalescent model to construct the phylogeny with 1,000,000 random quartets and 100 bootstraps.

### 1.7 Designation of species and subspecies

The genomic methods described above were used to determine the evolutionary relationships among the sampled populations. Our assumption is that evolutionary distances among populations provide support for designation of species and subspecies within the SDC.

For this study, we started with the accepted designation of speckled dace as a single species throughout its wide, geographically diverse range [3]. We then used the Unified Species Concept to evaluate evidence that there are multiple lineages within the accepted speckled dace subspecies that might be distinct enough to qualify as species [27]. We selected the Unified Species Concept because it provides flexibility in determining species, given that speckled dace hybridize readily with other cyprinid species, a common phenomenon among cyprinids.

Evidence needed to support likely species using genomics included (a) previous designation as a species based on conventional taxonomy, using morphological and meristic traits, (b) co-occurrence with other fishes endemic to a particular region, and (c) distribution limited to a geographically defined area with an underlying geology that indicates a high likelihood of long reproductive isolation. Sample sites were selected based on these criteria before the project started. Subspecies determination used the same criteria although we do not expect subspecies to be as differentiated from one another as species.

## Results

### 2.1 Sequencing, de novo RAD assembly, alignment

To assess the sequencing quality, we calculated the depth at 50 bp in each RAD contig. The mean individual coverage (i.e., the average coverage across all the contigs in one individual) was 7.69, with a maximum 24.88, a minimum of 2.50, and a standard deviation of 3.92 (S1 Fig). The final assembly contained 17,639 contigs, with a mean of contig length of 456.20, a maximum length of 788, and a minimum length of 89 (S3 material). After filtering individuals with sequencing and mapping quality, there were 175 individuals and 421,929 SNPs for further analyses (S1 Appendix).

### 2.2 Range-Wide

Across all speckled dace samples range-wide, the first two PCs explained 16.7% of the total variance (S2A Fig) and our samples were divided into three clusters (Fig 2A). Group One (upper right) consists of the Klamath speckled dace and Sacramento speckled dace subspecies as well as undesignated speckled dace collected from Warner Basin and Butte Lake. Group Two (upper left) is made up of the following designated subspecies: Amargosa, Long Valley, Owens, and Lahontan speckled dace. Group Three (lower middle) is composed of the Santa Ana speckled dace subspecies as well as undesignated speckled dace from the Bonneville Basin, Washington Coast, Columbia River, and Lower Colorado River. Speckled dace from the latter four regions are outside of California but serve to indicate that California populations sampled herein are distinct from populations in the rest of the range of speckled dace. We next used an admixture analysis of all the samples to complement our PCA analysis. The admixture analysis was run with K = 2-9 (S3 Fig). At K=3, members of each group in admixture analysis comprise Group One, Group Two and Group Three as indicated by PCA (Fig 2B). Furthermore, pairwise F_st_ calculated between subspecies varied from 0.16 (Owens and Amargosa speckled dace) to 0.68 (Amargosa and Santa Ana speckled dace) (S2 Table). Taken together, these results revealed that the subspecies in Moyle (2002) [1] have highly variable levels of genetic divergence and taxonomic revision may be warranted.

**Fig 2.**
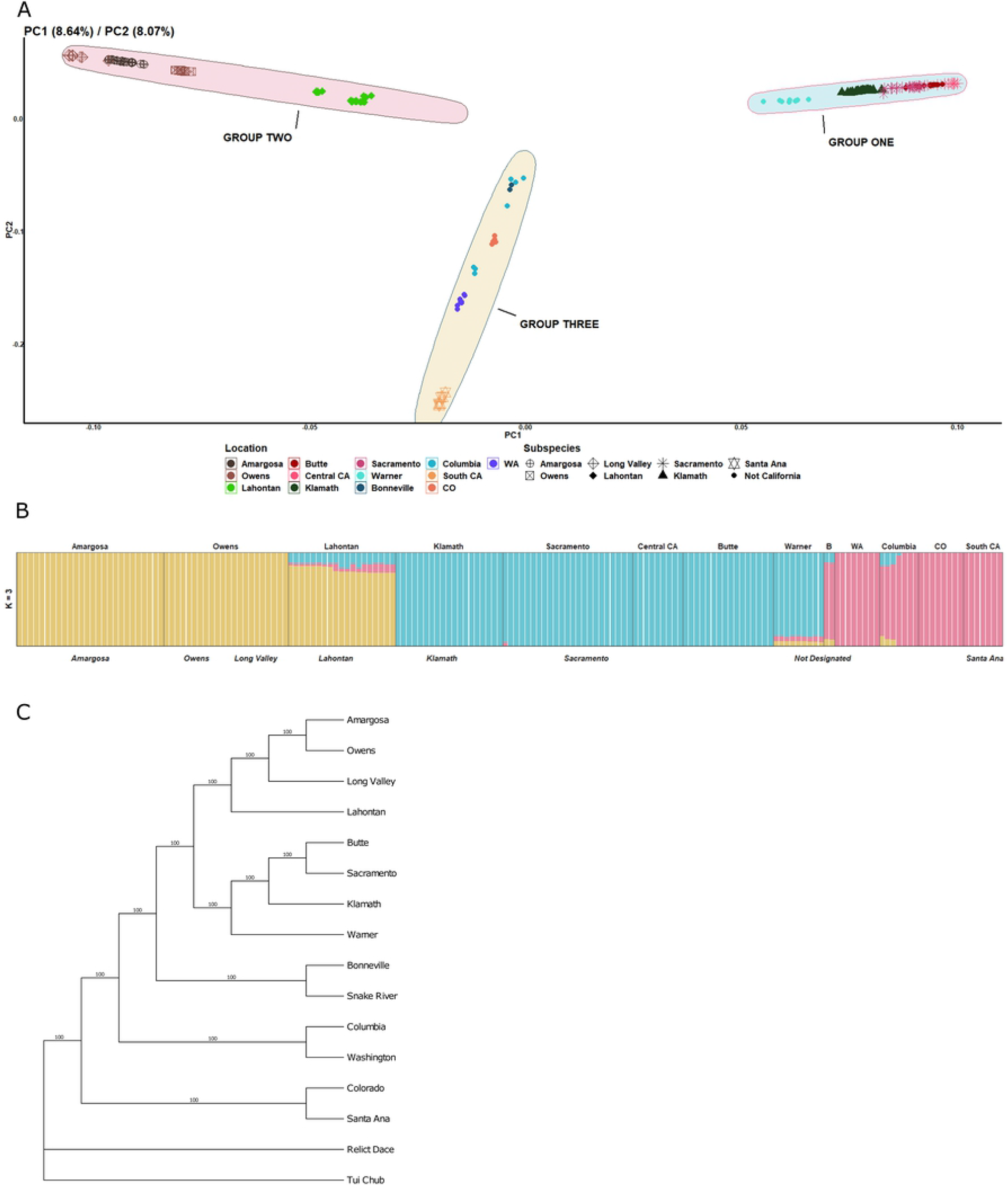
Range-wide speckled dace population structure. **A**. Principal Component analysis (PC) of all samples. Color and shape represent locations and subspecies designated in Moyle (2002), respectively. 16.71% genetic variation is explained in total (PC1 explains 8.64% variation while PC2 explains 8.07% variation). Three groups are distinguishable. Group One includes Sacramento speckled dace, Klamath speckled dace, and speckled dace collected from Butte Lake and Warner Basin. Group Two includes Amargosa, Long Valley, Owens, and Lahontan speckled dace subspecies. Group Three includes Santa Ana speckled dace and speckled dace from outside California which were collected from Washington Coast, Columbia River, Bonneville Basin, and Colorado River Basin. **B**. Admixture analysis of all samples when K = 3, which means we assumed the current populations are admixed by three populations in the past. The upper label represents the locations, and the lower label represents the subspecies designated in Moyle (2002). Washington, Colorado, and Bonneville are abbreviated as WA, CO, and B. PC analysis results are supported by results from Admixture analysis; the colors in three graphs therefore correspond. **C**. SVDQuartets results of the range-wide samples. Relict dace and tui chub were used as the outgroup. Speckled dace taxa designated in Moyle (2002) are split into three groups. Group One and Group Two are monophyletic and are the sister groups of each other, while Santa Ana speckled dace were clustered with Colorado Speckled dace and were the sister group of all the other speckled dace included in this study.

Our SVDQuartets range-wide phylogenetic analyses indicated that the speckled dace subspecies in California recognized by Moyle (2002) are mainly distributed into two monophyletic groups, with the exception Santa Ana speckled dace which is a distinct evolutionary lineage (Fig 2C). Similar to the results of PCA and admixture analyses, Sacramento, Klamath speckled dace and undesignated speckled dace collected from Butte Lake and Warner Basin belong to the same monophyletic group (Group One). Lahontan, Long Valley, Amargosa, and Owens speckled dace are located in another monophyletic group (Group Two), which is the sister group of Group One. Santa Ana speckled dace are clustered with speckled dace from the lower Colorado River drainages, and they are a sister lineage to (a) all other California speckled dace, (b) speckled dace from Bonneville and Snake River (separated from Columbia region because of admixture), and (c) speckled dace from Washington Coast and the Columbia River. Overall, the genetic structure/divergence of speckled dace in California is hierarchical rather than evenly distributed among the subspecies listed in Moyle (2002) (Fig 2).

### 2.2 Group One

Group One speckled dace include Klamath and Sacramento speckled dace subspecies plus speckled dace collected from Butte Lake and Warner Basin which were not designated in Moyle (2002). After our range-wide data set indicated Group One, we performed additional PC and admixture analyses using only Group One samples. For this PCA, the first three PCs explain the largest proportion of the genetic variation (S2B Fig). PC1 explains 7.59% genetic variation and split Warner Basin from speckled dace from the other regions. PC2 explains 4.69% genetic variation, and split Klamath from Sacramento speckled dace (Fig 3A). PC3 explains 3.73% genetic variation and separates the subregions within Sacramento speckled dace: speckled dace in Central Coast are separated from speckled dace in the Sacramento region (Pit River, Goose Lake, and Sacramento River) (Fig 3B). Speckled dace from Butte Lake cluster with Sacramento speckled dace in all the PCs, indicating genetic similarity. Admixture analysis support the results from PCA: Klamath speckled dace, Sacramento speckled dace, speckled dace collected from Warner Basin and Butte Lake were split gradually. More specifically, admixture analysis split speckled dace from Warner Basin when K = 2, and then Klamath and Sacramento speckled dace are distinct when K = 3. At K=4, Butte Lake are split from Sacramento speckled dace. Pairwise F_st_ analysis also support the results from PC and admixture analyses; the highest F_st_ values are found between Warner and the other locations (mean: 0.32) (S2A Table), whereas the F_st_ value between Klamath speckled dace and Sacramento speckled dace is 0.18. The F_st_ value between Sacramento speckled dace and Butte Lake speckled dace is only 0.086.

**Fig 3.**
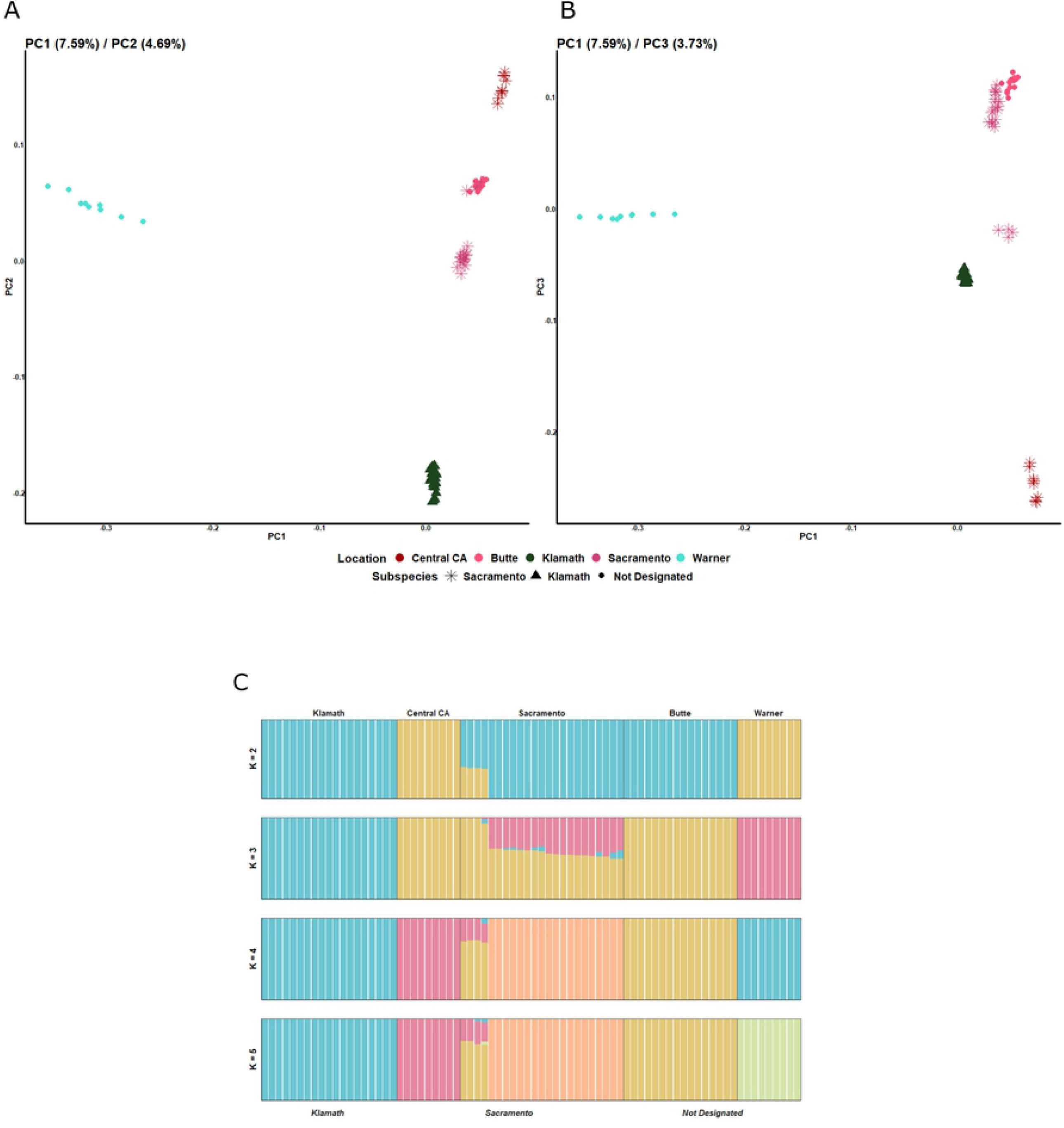
Sacramento-Klamath-Warner speckled dace population structure. **A**. Principal Component analysis of samples in group one; color and shape represent locations and subspecies designated in Moyle (2002), respectively. 12.28% of the genetic variation is explained by PC1(7.59%) and PC2 (4.69%). **B**. PC analysis of samples in group one when genetic variation is explained by PC1 (7.59%) and PC3(3.73%). C. Admixture analysis of samples in group one when K = 2, 3, 4, 5. The upper label represents locations, and the lower label represents subspecies in Moyle (2002).

The range-wide SVDQuartets analysis is concordant with the above results. The phylogeny placed, Sacramento, Klamath, Central California, Warner Basin and Butte Lake speckled dace into one clade (Bootstrap support =100). Within this clade, Sacramento and Butte Lake show great genetic similarity in PC and admixture analyses, and F_st_. Klamath speckled dace is the sister group to Sacramento speckled dace and Butte Lake, and accordingly Klamath was split from Sacramento speckled dace in PC and admixture analyses and F_st_. Warner speckled dace, which is the sister group of all the other groups, has the split in the smallest K value in admixture analysis, and based on pairwise Fst and PC analysis is the most differentiated from Sacramento and Klamath populations.

### 2.3 Group Two

Speckled dace clustered in Group Two include samples from three locations in the Death Valley region (Amargosa, Owens, and Long Valley) and Lahontan speckled dace. Each of these four locations are designated as a subspecies in Moyle (2002). To investigate the genetic structure within Group Two, we performed PC analysis, admixture analysis only on these samples. The first two PCs explain the largest proportion of the genetic variation (Fig S2C): PC1 explains 10.30% and PC2 explains 9.83% of the genetic variation among these samples. Amargosa and Owens speckled dace are very close to each other in both PCs; both PC1 and PC2 split Lahontan and Long Valley speckled dace from Owens and from Amargosa (Fig 4A). Admixture analysis supports the results of the PC analysis. Lahontan speckled dace are split from all the other speckled dace when K =2, and Long Valley is split from Owens and Amargosa when K = 3. At K=4 and K=5, we observed the local substructure in Amargosa speckled dace which is not discussed in this paper (Fig 4B). Although not as obvious as in Group One, F_st_ results support the PC and admixture analyses. The F_st_ between Owens and Amargosa speckled dace is 0.16, which was the smallest value in all the pairwise F_st_ values in Group Two; this is consistent with their close distance in PC analysis and differentiation at higher K values using admixture analyses. The F_st_ values between Long Valley-Owens and between Long Valley-Amargosa are 0.38 and 0.30, respectively, which is concordant with their separation in the PC analyses and early split in admixture analyses (S2B Table). However, Lahontan speckled dace, the first lineage to split in the admixture analysis, did not show the highest pairwise F_st_ values. Though the F_st_ value between Amargosa and Lahontan speckled dace is 0.33, this value was intermediate between F_st_ values for Lahontan-Owens and Lahontan-Long Valley. This presumably is the result of multiple evolutionary events such as hybridization between taxa or genetic drift in a dynamic landscape. The region has been active geologically during the Pleistocene with major filling and drying of lakes and a huge volcanic eruption that created the crater in Long Valley [1,3].

**Fig 4.**
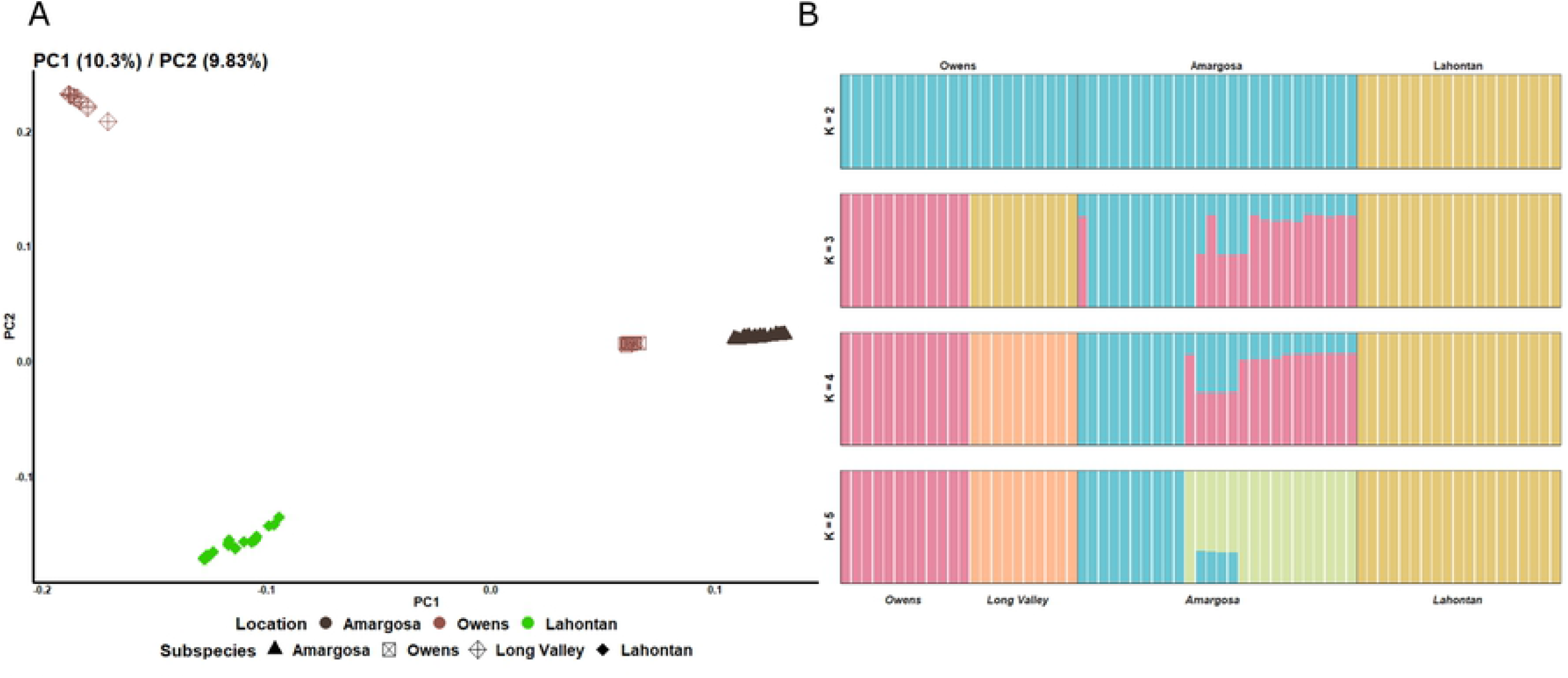
Death Valley speckled dace and Lahontan speckled dace population structure. **A**. Principal Component analysis of samples in Group Two; color and shape represent locations and subspecies in Moyle (2002), respectively. 20.13% of the genetic variation is explained by PC1(10.30%) and PC2 (9.83%). **B**. Admixture analysis of samples in Group Two when K = 2,3,4,5. The top labels are sample locations, and the bottom labels are subspecies designated in Moyle (2002).

The range-wide SVDQuartets analysis is concordant with PC and admixture analyses in Group 2. Lahontan speckled dace, which split at the beginning of admixture analysis, is the sister group of all Death Valley speckled dace. Owens and Amargosa, which show little genetic divergence in the PC and admixture analyses, are sister lineages in the SVDQuartets analysis. The position of Long Valley speckled dace in the SVDQuartets analysis is also supported by admixture analysis, where Long Valley splits after Lahontan but before Amargosa and Owens (Bootstrap support = 100). Although PC analyses and pairwise F_st_ indicate that Long Valley speckled dace is a separate lineage from Amargosa and Owens, this incongruence could be caused by overestimation of genetic divergence due to genetic drift in a small population under long isolation.

### 2.4 Group Three

The only California speckled dace sample in Group Three is the Santa Ana speckled dace, which clusters with non-California speckled dace (S1 Appendix). Group Three includes Santa Ana speckled dace, which is designated as a subspecies in Moyle (2002) and speckled dace collected from four non-California locations. To investigate the distinctiveness of Santa Ana speckled dace, we performed PC analysis, admixture analysis, and estimated pairwise F_st_ for sample collections in Group Three. PC and admixture analyses show that Santa Ana speckled dace are strikingly genetically different from non-California speckled dace in the group. In the PC analysis for Group Three, the largest proportion of the genetic variation is explained by PC1 and PC2 (S2D Fig). PC1 explains 21.5% and PC2 explains 12.9% genetic variation (Fig 5A). Strikingly, both PC1 and PC2 split Santa Ana from all other speckled dace lineages. Admixture analysis for Group Three was run from K = 2 to K = 5, and Santa Ana speckled dace split from the other locations such as Lower Colorado Basin, Bonneville, Columbia Basin, and Washington coast from K = 3 to K = 5 (Fig 5B). The F_st_ also show that Santa Ana speckled have high pairwise F_st_ values with all other California speckled dace and the non-California speckled dace (S2 Table).

**Fig 5.**
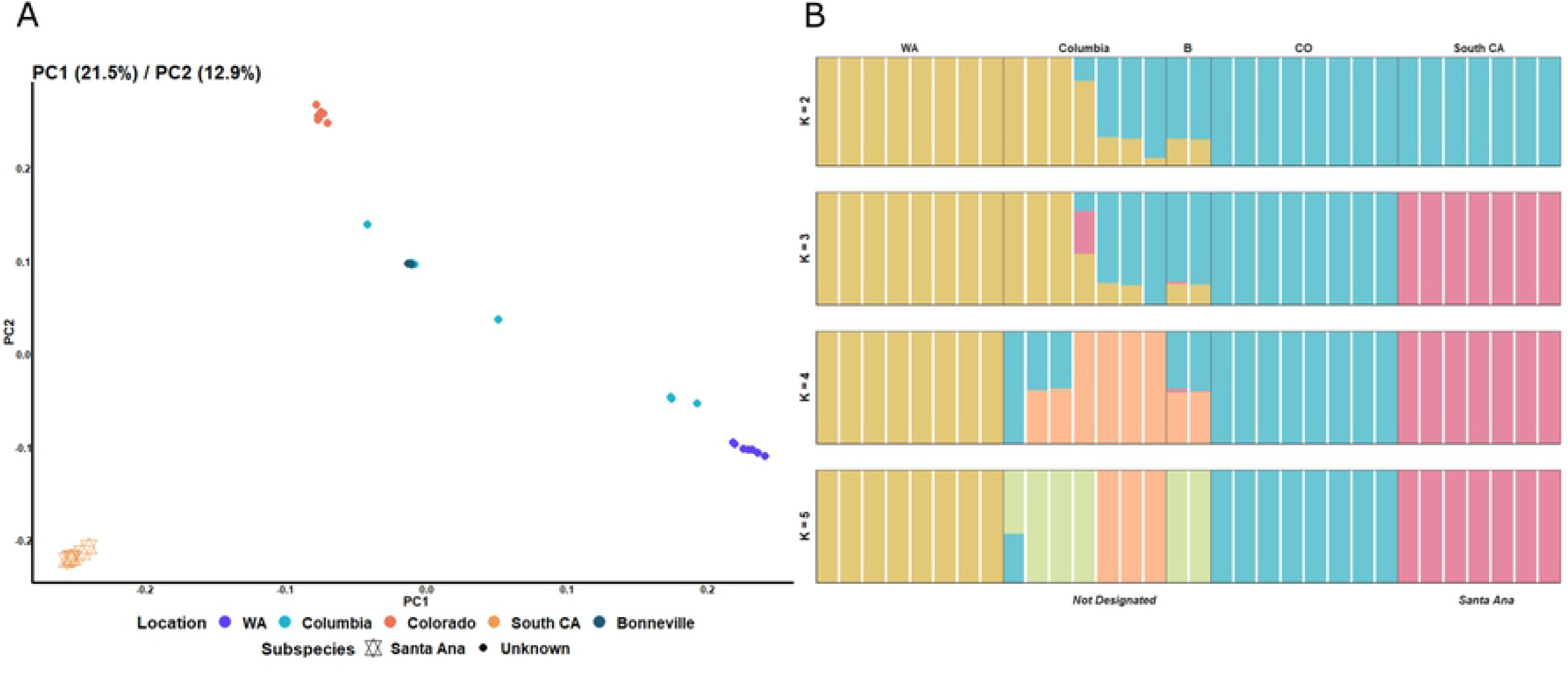
Non-California and Santa Ana speckled dace population structure. **A**. Principal component analysis of samples in Group Three. Color and shape represent locations and subspecies designated in Moyle (2002), respectively. 34.4% genetic variation is explained in total (PC1 explains 21.5% variation while PC2 explains 12.9% variation). **B**. Admixture analysis of samples in group three when K =2,3,4,5. The upper label represents the locations, and the lower label represents subspecies designated in Moyle (2002). Both PC and admixture analyses support Santa Ana speckled dace as distinct from non-California speckled dace but have a distant relationship to Colorado Basin dace.

The range-wide SVDQuartets analysis showed that Santa Ana speckled dace are clustered with samples collected from Colorado Basin (Bootstrap support = 100). The results of admixture analysis and PCA support the genetic affinity between Santa Ana and the Colorado Basin: Colorado speckled dace clusters with Santa Ana when K = 2 and Colorado speckled dace are the closest lineage to Santa Ana in the PCA. However, due to the limited number of samples of non-California speckled dace, we did not assess further the relationship between non-California speckled dace and Santa Ana speckled dace.

## Discussion

### 3.1. The speckled dace is a complex of multiple species and subspecies

Our genomic data analyses suggest that the speckled dace is not one species but rather a species complex with hierarchical evolutionary lineages, some of which may be designated as species and subspecies. In California, these lineages coincide with zoogeographic regions that are largely isolated from one another and that contain other endemic fishes, suggesting long isolation [1]. While some morphological and meristic differences exist among the lineages within the speckled dace, as discussed in the introduction, they may reflect local adaptations to diverse conditions rather than traits that allow species to be defined. Smith et al. (2017) [3] indicated that the lack of clear morphological differences was the result of frequent hybridization events that allowed gene flow among populations over wide areas. Because hybridization is common in cyprinid fish, we rely on pre-mating isolation as the basis for designating species and subspecies. We followed a combination of genetic and ecological differences to designate species and subspecies. Ecological differences and allopatric isolation ensure that genetic differences will accumulate. Thus, we hypothesize that a hybrid between individuals from two distinct lineages is expected to be poorly adapted to the ecological system and therefore have reduced fitness [28]. We therefore find it appropriate to label geographically isolated lineages with large genomic differences from other lineages as species and geographically isolated lineages with less genomic differentiation as subspecies or as distinct population segments [27]. A more comprehensive definition of species in the SDC complex will be presented in a separate paper that describes species and subspecies in California which are currently all under *R. osculus*.

### 3.2 Group One: Klamath, Sacramento, and Warner speckled dace together are a single species, with three subspecies

In all the analyses, Klamath speckled dace, Sacramento speckled dace, and undesignated speckled dace from Warner Basin are in Group One. Speckled dace in Group One have relatively low F_st_ values within them compared to F_st_ values between them and the other groups. For example, F_st_ values between Sacramento and Klamath speckled dace and speckled dace collected from Warner Basin are 0.18 and 0.30, but the F_st_ values between Sacramento speckled dace and Death Valley speckled dace (Amargosa, Owens, Long Valley) are 0.56, 0.52, 0.46 respectively. Samples from group one locations are also located in a monophyletic group in phylogeny. As a result, Klamath, Sacramento, and Warner Basin speckled dace should be considered a single species, the western speckled dace. In the genetic analysis of Group One, Warner is distinct from Klamath and Sacramento. The Warner Basin is also comparatively small, geographically well defined, and contains other endemic fishes. Therefore, we recognized Warner speckled dace as a subspecies under western speckled dace. Although Klamath and Sacramento speckled dace have less genetic divergence from each other than either does from Warner, the geographic basins in which each occurs are well defined and contain endemic fishes [1]. As a result, we also define Klamath speckled dace and Sacramento speckled dace each as a subspecies under western speckled dace. Speckled dace from the Central Coast of California (San Luis Obispo Creek, Santa Maria River, Monterey Bay drainages) are clearly part of Sacramento speckled dace lineage but show enough genetic differentiation that Central Coast speckled dace can be recognized as a separate DPS of Sacramento speckled dace (California coast speckled dace). Thus, Warner speckled dace, Klamath speckled dace, and Sacramento speckled dace are three subspecies, each under western speckled dace, of which only one of the three has been formally described, as *R. klamathensis* [29].

### 3.3 Group One: speckled dace from Butte Lake in Lassen Volcanic National Park is an introduced population

Butte Lake is located in Lassen Volcanic National Park and drains into the Lahontan basin, so speckled dace from Butte Lake were assumed to be genetically tied to Lahontan speckled dace. However, our analyses showed Butte Lake speckled dace to have much greater similarity to our Sacramento speckled dace samples than to Lahontan speckled dace in all the analyses. Therefore we classify speckled dace from Butte Lake as Sacramento speckled dace and hypothesize that the population in Butte Lake most likely represents a bait-bucket introduction. Butte Lake drains northward from Mount Lassen through Butte Creek (which has Sacramento speckled dace) and may have been connected at one time to the Eagle Lake watershed in the Lahontan Basin, although frequent lava flows have obscured drainage patterns. The three other fishes present in Buttle Lake, Tahoe sucker (*Catostomus tahoensis)*, Lahontan redside (*Richardsonius egregius)*, and tui chub (*Siphatales bicolor*) are Lahontan basin fishes, lending credence to the bait bucket hypothesis.

### 3.5. Group Two: Amargosa and Long Valley speckled dace are Subspecies of Lahontan speckled dace

In our phylogenetic analysis, Owens, Amargosa, Long Valley, and Lahontan speckled dace are in the same clade, which is in accordance with the results shown in range-wide PC and admixture analyses. In the genetic analysis for Group Two, Amargosa and Owens only showed show small genetic differences and a similar pairwise F_st_ value to the F_st_ value between Sacramento and Klamath speckled dace which we designate as separate subspecies. Smith et al. (2017) found that speckled dace from the Amargosa River shared haplotypes with speckled dace from Owens Valley. Dace from Oasis Valley, Nevada, headwaters of the Amargosa River in Death Valley and Ash Meadows (Bradford Spring), are sister lineages of Owens Valley speckled dace. Mussman et al. (2020) also showed that speckled dace from Owens and Amargosa watersheds had minimal genetic variance and that admixture exists between speckled dace from these two regions. Unlike the situation for Klamath and Sacramento speckled dace, the Owens River and Amargosa River watersheds are internal drainages that were connected via a chain of large lakes during extended wet periods in the late Pleistocene, 16-18 thousand years ago [30-31]. Given the results of our analyses and their recent geographic separation and isolation, we place speckled dace from Amargosa and Owens Rivers in the same subspecies, named Amargosa speckled dace. However, although Long Valley speckled dace are in the same geographic basin as the Owens watershed, Long Valley speckled dace are genetically distinct from Owens and Amargosa speckled dace. This was probably the result of genetic drift due to isolation of small dace populations by a large volcanic eruption that created Long Valley and isolated the Owens Valley from the Lahontan Basin. Therefore, we consider Long Valley speckled dace to be a subspecies of Lahontan speckled dace. All other populations of speckled dace in the Death Valley region (e.g., Owens Valley, Ash Meadows, Amargosa River, Oasis Valley) are best regarded together as one subspecies of Lahontan speckled dace, with each designated as a Distinct Population Segment (as in [15]). Overall, our analyses points to Amargosa speckled dace and Long Valley speckled dace each as being subspecies of Lahontan speckled dace, which is discussed section 3.5.

### 3.5. Group Two: Lahontan speckled dace are a species with three subspecies in California

Speckled dace from the Walker River, Humboldt River, Eastern Sierra Nevada streams, and Death Valley system streams are supported as one lineage, the Lahontan speckled dace, which is a widely recognized taxon, as *R. o. robustus* [1,33,34]. The Owens, Amargosa and Long Valley populations form lineages that have diverged from Lahontan speckled dace and could arguably be recognized as a full species (Fig 2B). Although Lahontan speckled dace split at K=2 in the admixture analysis for Group Two, F_st_ values between Lahontan-Owens and Lahontan-Long Valley speckled dace are somewhat small: 0.25 and 0.26, respectively, and even lower than the F_st_ between Long Valley-Owens speckled dace. This suggests that either a hybridization event took place between Lahontan-Owens creating Long Valley speckled dace, or that Lahontan and Long Valley share an early evolutionary history and were separated by geologic change. Ancestral Lahontan speckled dace were likely present in the Owens region prior to a massive volcanic eruption that separated the Owens Valley from the Lahontan basin about 760,000 years ago [35]. The presence of Owens tui chub (*Siphatales bicolor snyderi*) and Owens sucker (*Catostomus fumeiventris*) support this concept because the closest relatives of both taxa are in the Lahontan basin [1]. Therefore, we consider Lahontan speckled dace to have two subspecies in California: Amargosa speckled dace and Long Valley speckled dace. However, the presence of other *R. osculus* subspecies, some described, in the Lahontan Basin indicates that additional subspecies will likely eventually be added to the list [34].

### 3.6 Group Three: Santa Ana speckled dace is a full species

Our range-wide analyses reveal that Santa Ana speckled dace are strikingly different from all other speckled dace collected in California. Santa Ana speckled dace share more genetic similarities with speckled dace from Lower Colorado Basin, Bonneville, Washington Coast, and the Columbia River than with other dace in California. Due to the small number of samples, the genetic diversity within the Colorado Basin, Washington Coast streams and the Columbia River are not discussed in this paper. The evolutionary history of Santa Ana speckled dace can be linked most closely with speckled dace in the Lower Colorado Basin because they did not split from each other in the admixture analysis with all the samples from K = 3 to K = 8 (S3 Fig). In Smith et al. 2017, speckled dace collected from Colorado Basin and speckled dace collected from Los Angeles Basin are sister lineages in the Colorado Group with a relatively weak bootstrapping support in the MtDNA phylogeny. In the Fst values, we find Santa Ana speckled dace has high genetic divergence from both California and non-California speckled dace. The lower bootstrapping in Smith et al. (2017) [3] is likely caused by high genetic divergence and relatively few diversity in MtDNA.

In our study, we clarify the genetic distinctness of Santa Ana speckled dace. All analyses show that Santa Ana speckled dace have remarkably high genetic differences from the other subbspecies in Moyle (2002) [1] and the other speckled dace in Group Three (S2A and S2B Tables). Due to the distinct genetic structure separating Santa Ana from other sampled California speckled dace in addition to those from the Columbia and Colorado river basins, Santa Ana speckled dace clearly merit full species recognition. This same basic conclusion was reached by Cornelius (1969) [32] who conducted a detailed study of the morphometrics and meristics of Santa Ana speckled dace, as well as of dace from neighboring streams (Sacramento basin), the Virgin River (Lower Colorado basin), and Lake Tahoe (Lahontan basin). His study was the first to link the origins of Santa Ana dace to the lower Colorado River basin. Using genomics, Mussman (2020) [15] came to the conclusion: that Santa Ana speckled dace are very different and are linked to a Colorado River clade.

### 3.7 Conservation Implications

We found that genetic divergence in speckled dace is concordant with geographical regions and has a hierarchical structure: the populations across geographical regions are genetically divergent in different levels, depending on time and degree of isolation from other speckled dace populations. If we view the SDC as a single widespread species, it would not be considered as a species that needs conservation because of its wide distribution and large population size. Species and subspecies in different geographical regions face different environmental problems. We can combine our knowledge of genetic divergence with that of ecosystem status and characteristics to design distinct conservation management and policy strategies for different populations of speckled dace. Such information can also set help priorities for conservation: which lineages of speckled dace need conservation attention now in order to protect genetic diversity.

## 4. Conclusion

Based on the genetic analyses, we found that the relationships within the seven subspecies designated in Moyle (2002) [1] are hierarchical. In other words, they are genetically divergent at different levels as opposed to having relatively uniform relatedness as might be expected for a single taxonomic level. This result supports merging lineages with relatively small genetic differences into single subspecies and treating the most genetically distinct lineages as species. More specifically, our genetic analyses place all California populations into three species: western speckled dace, Lahontan speckled dace, and Santa Ana speckled dace. Western speckled dace contains, as subspecies, Warner speckled dace, Sacramento speckled dace, and Klamath speckled dace. Lahontan speckled dace is a species that is widely distributed in the Great Basin but that also encompasses two lineages of speckled dace from the Death Valley region: Long Valley speckled dace and Amargosa speckled dace. Santa Ana speckled dace is also a full species showing extreme genetic divergence from all other speckled dace. The focus of this paper is speckled dace in California so how our findings relate to speckled dace outside of California remains unclear. However, it seems likely that there are non-California lineages that can also be designated (or redesignated) as full species when genomic methods are applied to uncover cryptic species. These taxonomic issues will be further explored in a paper devoted solely to taxonomy of the speckled dace complex in California.

## Acknowledgements

The authors thank Amber Manfree for making the maps (Fig 1) for this project as well as Scot Lucas (Klamath River), Mollie Ogaz (Sacramento system), and Steve Parmenter (Owens Valley, Lahontan) for providing samples for this analysis.

## Supporting Information

**S1 Fig. Contig Depth**. The distribution of mean depth of all the contigs at 50 bp in each individual.

**S2 Fig. Genetic variation explained by each PC. A**. The percentage of genetic variation explained by first 30 PCs for all samples PCA. **B**. The percentage of genetic variation explained by first 30 PCs for PCA for California speckled dace, Warner speckled dace and speckled dace in Butte Lake (Group one). **C**. The percentage of genetic variation explained by first 30 PCs for PCA for Death Valley speckled dace and Lahontan speckled dace (Group two). **D**. The percentage of genetic variation explained by first 30 PCs for PCA for Non-California and Santa Ana speckled dace (Group three).

**S3 Fig. Admixture analysis of all the samples from K = 2-9**. K refers to the number of the ancestral populations that the current populations are admixed from. Each color represents one of the ancestral populations. Upper and lower x-axis refers to locations and subspecies, respectively.

**S1 Appendix. Information for all samples included in the analysis**. It includes the location and GPS coordinates and number of the samples after the sequencing and alignment qualifying filtering.

**S1 Table. Number of SNPs in the analyses for each group**. It counts number of SNPs that are genotype called in PCA and admixture analyses for investigating the population structures in group 1-3.

**S2 Table. Pairwise F**_**st**_ **for speckled dace**. The smaller the number the closer the evolutionary relationship between the two populations. **A**. Pairwise F_st_ of California samples and samples that share the same node with California samples. **B**. Pairwise F_st_ between non-California samples (Washington, Columbia, Colorado, Washington) and Santa Ana speckled dace.

**S1 material. Individuals selected for reference genome**. The file names of the filtered BAM files of 8 speckled dace collected from Walker River, which are selected to generate reference genome.

**S2 material. Outgroup Sequence**. It includes the sequence of tui chub and relict dace used as outgroup for phylogenetic analysis.

**S3 material. The list of Contigs in the reference genome**. This file contains all the contigs included in the reference genome with the average, minimum, and maximum length.

